# MirMachine 2: a scalable, evolutionarily informed pipeline for microRNA annotation and comparative genomics across thousands of animal genomes

**DOI:** 10.64898/2026.05.19.726197

**Authors:** Vanessa M. Paynter, Sinan U. Umu, Jack A S Tierney, Francesca Floriana Tricomi, Leanne Haggerty, Bastian Fromm

## Abstract

Genome sequencing is rapidly outpacing the annotation of conserved regulatory elements, limiting the evolutionary and comparative insights that can be extracted from expanding genome collections. MicroRNAs are among the most conserved and phylogenetically informative genes, yet automated annotation has remained difficult to scale while preserving evolutionary interpretability. Here we present MirMachine 2, an evolutionarily informed framework that combines curated reference models, lineage-aware scoring, and adaptive filtering to enable robust genome-wide microRNA annotation at scale. Applying this to thousands of animal genomes reveals that many apparent absences of conserved microRNAs reflect methodological bias rather than biological loss, particularly in underrepresented lineages. By enabling consistent and interpretable comparison of microRNA complements across large datasets, MirMachine 2 establishes scalable microRNA annotation as a practical foundation for genome-scale evolutionary and comparative genomics.

## INTRODUCTION

MicroRNAs are evolutionarily conserved, short non-coding RNAs that function as post-transcriptional regulators of gene expression. They act as key regulators in vital biological processes including development, cell proliferation and apoptosis^1,2^ by inducing mRNA degradation or translational repression^3,4^. Owing to their high conservation and near hierarchical evolution, they preserve reliable phylogenetic signals^5–8^. Given their relevance in evolutionary^9–13^ and clinical studies^14,15^, accurate annotation of microRNA references is critical for meaningful downstream biological interpretation^16^. Gaps or misannotated elements can compromise phylogenetic reconstructions, bias comparative analysis of regulatory networks and mislead functional inference in species^17,18,19^ . Moreover, as genome sequencing efforts accelerate with over 23000 reference genomes currently available on NCBI, there is a pressing need for automated evolutionarily informed annotations that enable robust cross species comparisons providing a foundation for both evolutionary and translational research^20^.

Accurate annotation of microRNAs typically require high quality genomes, transcriptomic evidence, long trained expertise and manual curation making it a slow and resource intensive process. As a result, manual curation efforts, such as the 114 manually annotated species in our database MirGeneDB^21^, are unable to keep pace with the rapid influx of available genome assemblies. The development and introduction of our microRNA family covariance model (CM) and machine learning (ML) based tool MirMachine^22^ has helped address this limitation by enabling the automated and accurate annotation of conserved microRNAs, directly from genomes. In its initial deployment, MirMachine was successfully applied across diverse animal species, including extinct species where small RNA sequencing remains challenging. The study further exploited the accumulation of microRNAs through discrete evolutionary events to devise a simple microRNA scoring system as a quantifier for the completeness of conserved microRNA families. In this context we demonstrated MirMachine as an automated annotation tool that is robust and flexible while eliminating the dependence on smallRNA sequences.

However, MirMachine 1 was initially constrained in several respects that limited its use as a general framework for large-scale comparative genomics. For instance, its performance was only reported on a dataset of 88 eutherian genomes, leaving scalability across large and diverse data uncertain. And although the CMs were trained on the full complement of 75 different species available on MirGeneDB at the time, uneven taxonomic representation and gaps in available animal phyla led to reduced performance in certain invertebrate groups. Early implementations also lacked configurable parameters to accommodate for structural variability, and atypical features found in giant microRNA hairpins^23^. Given the relative heterogeneity of microRNA architecture across the tree of life, and deviations from “canonical” families, limitations in tunable features reduced the sensitivity for detecting divergent microRNAs. This substantially limited sensitivity for microRNAs deviating from representative models of respective families. In addition, a proposed scoring feature was exploratory in nature, and thus was neither benchmarked against established annotation metrics like BUSCO^24^, nor automatically implemented into the pipeline. Together, important questions regarding scalability, sensitivity, comparative performance, and standardised reporting remained unresolved and warranted further methodological refinement.

## DESIGN

In this study, we present MirMachine 2 and demonstrate the following methodological advancements: i) scalability to thousands of genomes, ii) implementation and benchmarking of the microRNA score to other established genome metrics, iii) lineage aware pre-filtering of results and iv) showcase on 785 lepidopteran genomes the performance of MirMachine 2 for biological insights.

To evaluate scalability MirMachine 2 was applied to the complete Ensembl genome assembly dataset comprising 3019 metazoan genomes at the time^25^. In support of this expansion, microRNA family covariance models were retrained on the recently updated complements on MirGeneDB 3.0 consisting of 114 species providing a larger representative dataset thus increasing taxonomic coverage. MirMachine 2 integrates the microRNA score as an automated feature informed by evolutionarily known microRNA gains and losses^21^. We further explore its relationship to widely used genome contiguity and completeness metrics such a N50 and BUSCO and find a strong congruence among them. Thereby, revealing the potential for the microRNA score as a complementary indicator of assembly quality alongside the microRNA complement completeness.

Additionally, MirMachine 2 enables manual scrutiny of predictions from underrepresented taxa that behave unexpectedly despite retraining and confidence levels. For such cases, we propose a simple decision-tree that leverages paralog expectations, seed conservation and structural integrity. To support this, we have integrated seed sequence identification with conserved “star” seed motifs explicitly flagged in the output alongside confidence of each prediction. This facilitates direct comparison of annotations to help distinguish potential biological phenomena such as possible lineage specific losses and pseudogenes from annotation artefacts^22^. We also offer tunable parameters including an e-value for threshold stringency (-e) and extended precursor lengths (-long hairpin). Thereby improving sensitivity to structural and sequence variability across microRNA families. Collectively, these improvements encourage informed inspection of outliers rather than automatic exclusion based on rigid cutoffs particularly in under-represented taxa. Our results highlight the microRNA score and seed sequence conservation as robust parameters to assess complement completeness and incur possible biological implications where microRNA scores deviate from expected.

Altogether our study demonstrates MirMachine 2 as a computationally efficient, high throughput annotation platform enabling reproducible comparative analysis through an evolutionarily aware analytical framework capable of revealing lineage-specific signals.

## RESULTS

### microRNA complements of the complete Ensembl dataset

Following the updates addressing the conceptual limitations of MirMachine 1, we evaluated the newest version for its performance at genome scale. Applying MirMachine 2 to the entire genome catalogue available in Ensembl, 3019 genomes across 1938 distinct species^25^ were annotated. Encompassing a broad list of animal genomes, this dataset serves as an excellent source to demonstrate the high scalability of MirMachine 2. The resulting comprehensive set of annotations reveal lineage specific patterns through the presence and frequency of conserved microRNA families (Figure 1.A, 1.D).

**Figure 1.**
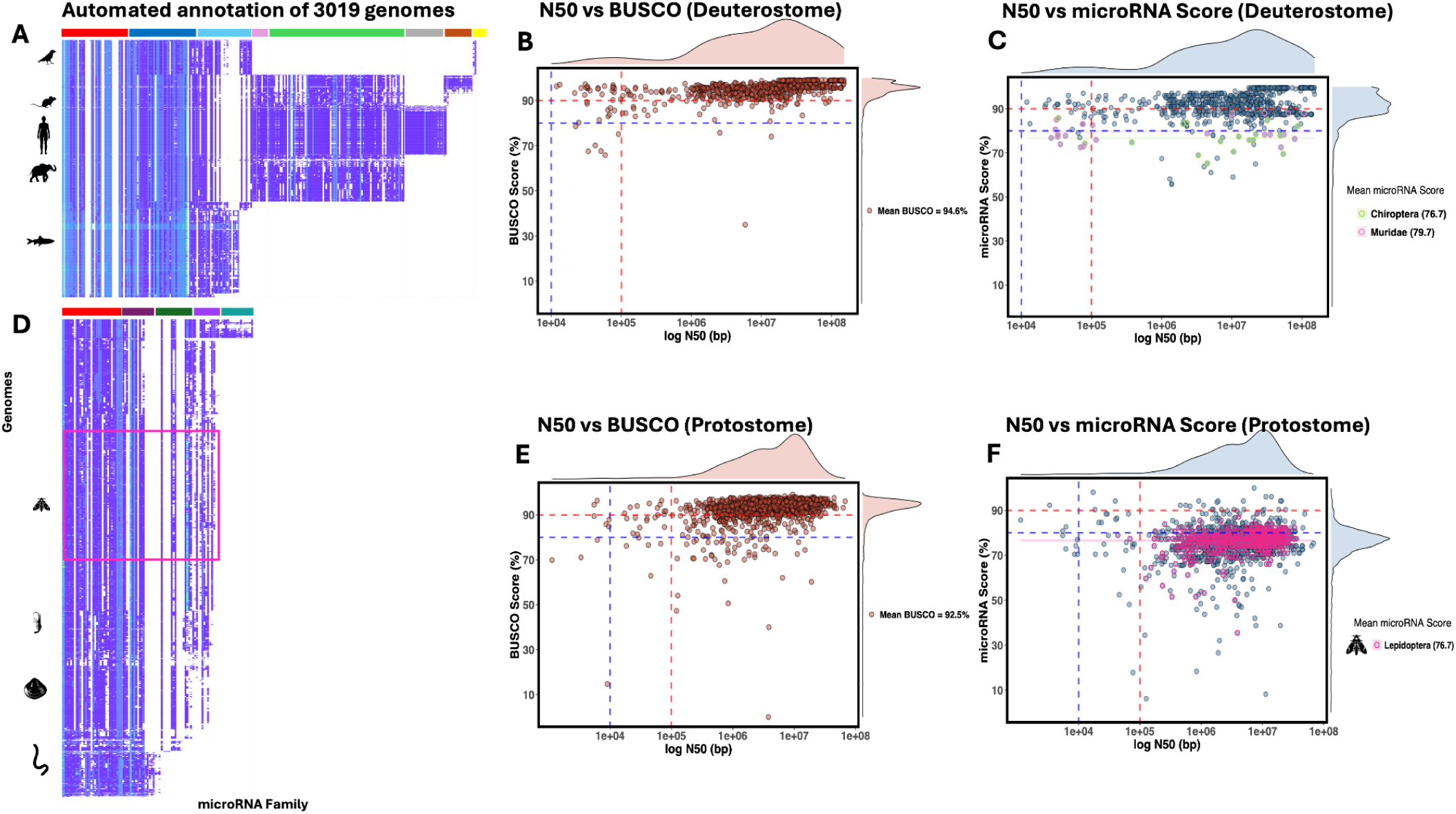
Conserved microRNA landscape across 3019 Metazoan genomes. A) Banner plot summarizing the presence, absence and frequency of conserved microRNA families across the complete vertebrate genome dataset from Ensembl. Colour bars indicate node specific microRNA family blocks as follows - Red: Bilateria, Blue: Vertebrata, Light Blue: Gnathostomata, Pink: Theria, Green: Eutheria, Grey: Catarrhini, Brown: Muridae, Yellow: Aves. B) Comparison of BUSCO completeness and assembly contiguity (N50) across deuterostome genomes. C) Comparison of microRNA scores and N50 across deuterostome genomes, highlighting outliers. D) Banner plot summarizing the presence, absence and frequency of conserved microRNA families across the complete protostome genome dataset from Ensembl. Highlighted are the lepidoptera genomes. Colour bars indicate node specific microRNA family blocks as follows - Red: Bilateria, Purple: Protostomia, Dark Green: Arthropoda, Lavender: Neoptera, Teal: Drosophila. E) Comparison of BUSCO completeness and assembly contiguity (N50) across protostome genomes. F) Comparison of microRNA scores and N50 across protostome genomes, where most genomes underperform unexpectedly (Lepidoptera genomes are highlighted in pink).

The predicted microRNA complements reveal strong clade specific conservation patterns consistent with established evolutionary stability of core microRNAs^17,18^. The banner plot illustrates broad contiguity while gaps signify the absence of expected conserved families, indicating either a technical drop out, known secondary absences in specific lineages (such as the loss of the chordate family MIR-4057 in vertebrates), or previously uncharacterised true biological loss^26–28^.

To better distinguish between these possibilities, we applied the newly included microRNA scores as an initial quantitative discriminator. The score is calculated as the number of conserved microRNA families detected relative to the number expected given the phylogenetic position of the species (a new parameter of MirMachine). We next compared the runtime per genome of the microRNA score with BUSCO score^24^ using a subset of mammal genomes. Here, MirMachine exhibits faster performance, completing analysis on average 4.2 times faster than BUSCO, with a proportionally lower processor load (Supplementary Figure 1).

**Supplementary Figure 1.**
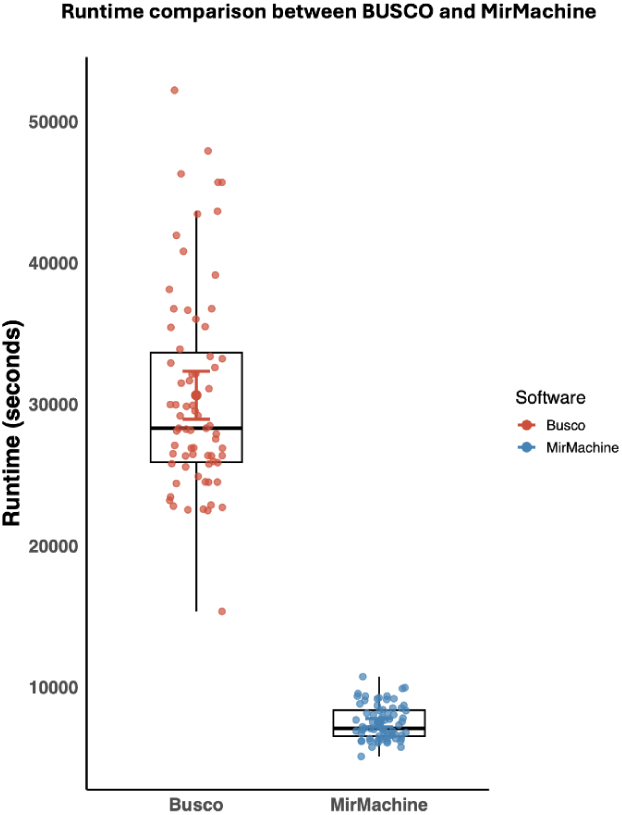
Comparison of runtime performance between BUSCO (mammals) and microRNA score. Mean run time across 71 mammal genomes with 95% confidence intervals.

### Performance and applicability of the scoring feature using Ensembl dataset

To assess the utility of the scoring feature for large scale comparative analysis, the microRNA scores of 3019 genomes were assessed in relation to assembly quality using N50 values that correlates with genome contiguity^29^. In parallel BUSCO completeness was assessed in relation to N50 as a comparison for benchmarking. As proposed by us earlier, this analysis showed that for genomes with high assembly quality typically considered over an N50 threshold of 1Mb, both scores remain consistently high. However, BUSCO scores tend to overperform in a plateau fashion likely due to its broader gene set. Notably, cnidarians and sponges displayed consistently low microRNA scores which can be attributed to their comparably small microRNA complement, resulting in unreliable scoring for phylogenetic groups where the complements may be small (Supplementary Table 1). Consequently, for downstream analysis these groups were omitted from the dataset to avoid skewing the analysis as they are not biologically informative to the scope of the current broader study.

The microRNA scores observed in deuterostome genomes (Figure 1.B, 1.C) reveal an expected high performance with a median score of 92.07, and only a few outliers such as those of bat and rodent genomes^22,30^. These outliers depict an uncharacteristically low score despite their high genome quality and well represented microRNA annotations. This finding follows a trend of lower microRNA scores as seen in Umu et al.^22^, and of those recently observed in rodents^30^. Contrastingly, a large proportion of protostome genomes revealed a shift in overall performance, with the median microRNA score at 77.04 (Figure 1.E, 1.F). A difference of over 10% between the BUSCO and microRNA score is significant given the difference in scale of the two measurements (Supplementary Figure 2). BUSCO assesses single copy orthologs (9226 genes for mammals using BUSCOv4), whereas the microRNA score focuses on a smaller functionally critical set of conserved microRNA families. Thus, it is expected that lower scoring for microRNA suggests either technical issues arising from incomplete genome assemblies or genuine biological signals due to loss of microRNAs in assemblies with higher quality.

To investigate the under-performance observed from high quality protostome genomes aggregating at lower microRNA scores, the BUSCO and microRNA scores were contrasted in a node based framework (Supplementary Figure 2). This revealed a phylum specific score reduction, most prominent in Neoptera, Diptera, Endopterygota and Lepidoptera genomes, despite a high N50.

**Supplementary Figure 2.**
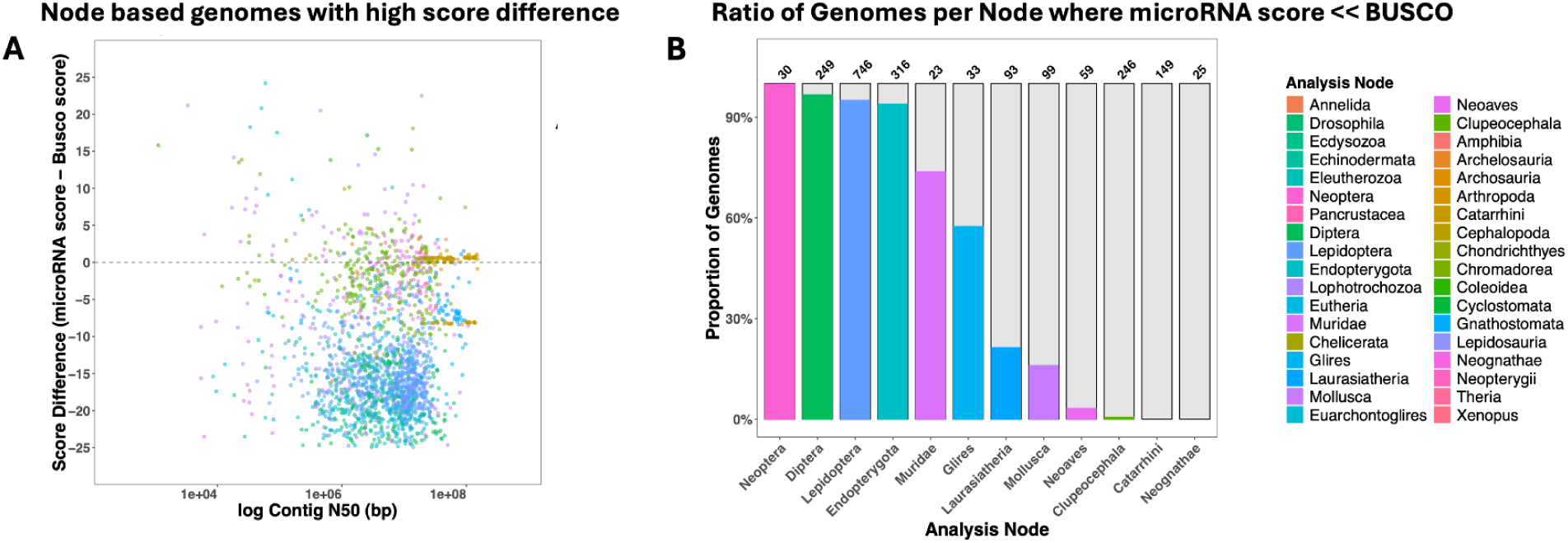
Phylogeny based distribution of microRNA score performance across genomes. A) Scatter plot contrasting the difference in score in relation to N50, coloured by phylogenetic node. B) Proportions of genomes exhibiting low microRNA scores summarized by nodes in taxa with >20 representative genomes are shown. Total number of genomes are indicated at the top.

Due to their large representation in the dataset, lepidoptera genomes were used to further examine patterns that reflect a broader trend for all insects. Although microRNAs can exhibit secondary mosaic losses^6,17^ (or reductions in parasitic groups^26–28^), they are uncommon and insufficient to explain the scale of observed reductions here. A likely contributing factor is the limited representation of these lineages in the training data, as a consequence of limited microRNA annotations for invertebrates. Initial models for MirMachine 1 were built on 1 species of Lepidoptera (Figure 2.B), represented by the annotation for *Heliconius melpomene* reducing the sensitivity for detecting variable families in these clades. We therefore examined the apparent absences in predictions to disentangle the effects of incomplete annotations and technical issues from genuine lineage specific loss, as addressed in the next section.

**Figure 2.**
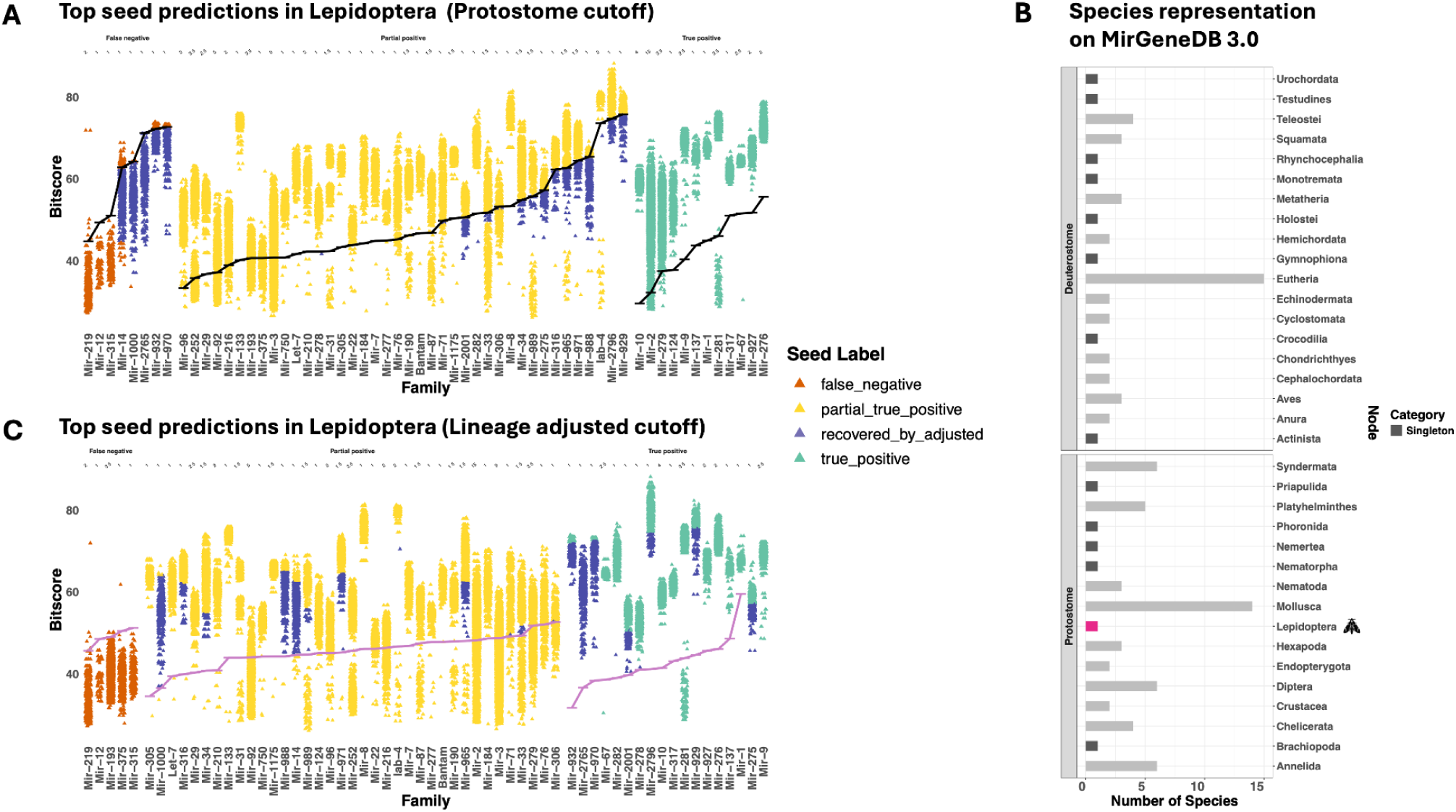
Lineage informed recovery of Lepidoptera specific microRNA families. A) Distribution of seed based predictions for all conserved microRNA families found in Lepidoptera, categorised by the proportion of predictions scoring above the default protostome threshold (black line) used by MirMachine. Purple points denote putative false negatives subsequently recovered through threshold adjustment. B) Summary table with number of species specific curated microRNA complements across MirGeneDB 3.0. C) Updated seed based predictions following the application of adjusted threshold (pink line) illustrating improved recovery.

### Exploring variation in scores/conserved complement predictions across phyla

Despite adequate genomic representation (716 genomes), the low scores in Lepidoptera raise the question of whether the absences can be explained as a genuine biological signal, or limitations in prediction thresholds as a consequence of insufficient species representation (Figure 2.B). Although the representative for Lepidoptera on MirGeneDB, has the whole complement annotated, it may not be sufficiently capturing possible variation across the node.

We observe the highest microRNA score of 88.7 in *Heliconius sara* followed by *Dryas iulia moderata* at 87.0 as opposed to a 100 for *Homo sapiens* genomes from the deuterostome dataset. We notice predictions containing conserved seed sequences usually drop out from the “filtered” output (high confidence predictions), suggesting filtering rather than an absence of expected families. Further implying that predictions may not be efficiently captured within the filtering system. This poses a risk of obscuring genuine biological signals where present, underscoring the importance of prediction accuracy in annotations.

To infer detection accuracy of true positives across families in underrepresented taxa a classification strategy was established. As an alternative to manual annotation and retraining, we used seed presence alongside paralog expectations as parameters to estimate detection accuracy. Expected seed counts were inferred from manual annotations of phylogenetically related species like *Heliconius melpomene* and *Drosophila melanogaster*. Top scoring seed predictions were then screened based on the average paralog expectation per family. We then classified the 62 expected conserved families for Lepidoptera according to the proportion of genomes in which their top performing seed predictions fell below the protostome threshold. Resulting in a gradient of detection accuracy as explained further.

The cutoffs used for filtering appear stringent, as an overall 21 families that display predictions containing seed conservation falls below the threshold resulting in the drop out observed with lepidopteran genomes. Upon binning (Figure 2.A, 2.C), 12 families were found to be “strict positive” with 0 failures across genomes. In this bin, families such as MIR-2, MIR-279 and MIR-281 occasionally showed seeds below the threshold, but given their high paralog numbers (eg, MIR-2 with 15 expected paralogs) these cases were rare and did not alter the overall classification in accordance to the binning strategy. Further, 40 families fall into the intermediate category where drop out of high performing predictions result in failure rates ranging from 0.1% to 78%. Here, most families (25) had a low dropout rate at 10%, and only 4 families (MIR-275, MIR-306, MIR-375 and MIR-988) showed a failure of above 50%. These intermediate cases highlight how stringent thresholds disproportionately affect families with moderate paralog numbers leading to systematic underreporting.

Finally, 8 families were binned as high false negatives since recovery of predictions was very low. The drop out of these families explain the lower microRNA scores from the lepidopteran genomes. Among these, MIR-14, MIR-1000, MIR-2765, MIR-932, MIR-970 appear “recoverable” (Figure 2.A, purple data points) owing to a clear demarcation between predictions with seed and non-seed sequences, emphasising the technical constraint in prediction accuracy for this taxa. This implies that the threshold can be further adjusted in a species specific manner as opposed to a blanket protostome cutoff for Lepidoptera.

Taken together, these results highlight that the protostome thresholds were overly conservative for Lepidoptera, leading to systematic under-reporting of microRNA families despite the evidence of conserved seeds in raw predictions. Consequently, families with higher paralog diversity and moderate sequence variation are unevenly affected. Predictions failing to fulfill the threshold requirements set by the training models can be attributed to the failure to capture enough variability due to the underrepresentation of lepidopteran microRNA complements in MirGeneDB. This limitation is apparent in later analysis comparing the annotation of nine economically relevant species as representatives of the clade, depicting fewer families retained in the final filtered output (Figure 3.A, 3.B). Inadvertently, such bias arises as sequencing efforts of invertebrates are uneven as previously surveyed^31^, ultimately creating a disparity in available microRNA annotations. More broadly, we demonstrate that fixed universal cutoffs may be too rigid to accommodate phylogenetic diversity.

**Figure 3.**
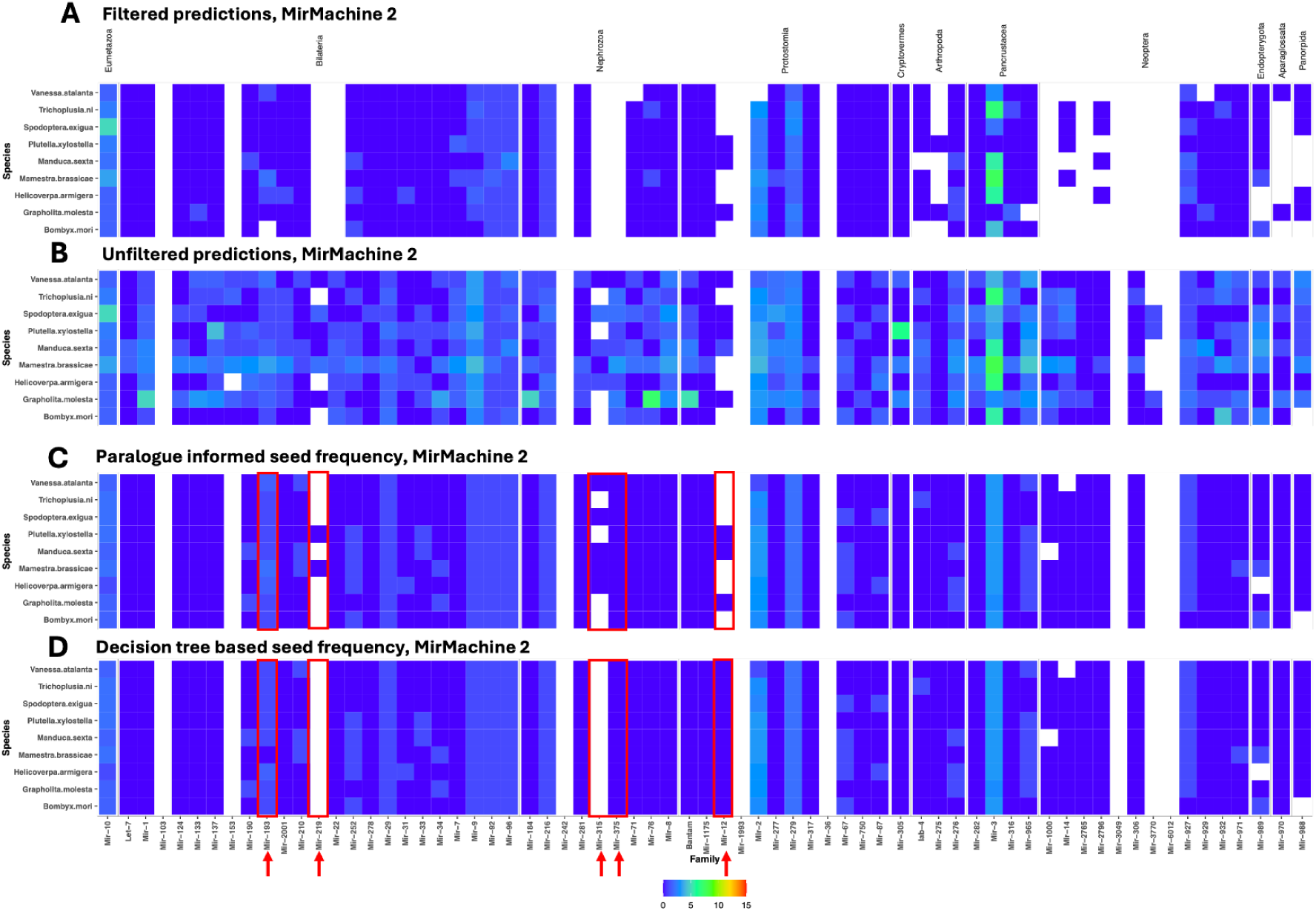
Comparison of microRNA prediction filtering based on the proposed decision-tree framework. (A) Filtered predictions generated by MirMachine 2. B) Unfiltered predictions generated by MirMachine 2. C) Predictions containing conserved seed motifs generated by MirMachine 2. D) Refined predictions highlighting the five low performing families under the decision tree.

Especially useful in cases where microRNA complements are incomplete as exemplified for the lepidopterans in this study. This inclusion of seed notations facilitates manual inspection of predictions particularly for families that fail to meet bitscore thresholds. In addition, star seed sequences are also highlighted reflecting the predominant seed associated with each microRNA family. Making it especially beneficial in analysing annotations of clades that are under-represented in MirGeneDB, as it allows for the closer inspection of otherwise filtered out predictions.

### Finding an optimal species specific threshold

The significant drop out of predictions as depicted after binning in figure (Figure 2.A) underscored the benefit in altering the cutoff thresholds to reduce the likelihood of false negatives (true but being left out). Thus, for improved recovery of true positives and to address the possible loss of families, a species specific threshold was redefined based on the bitscores of non-seed predictions as mentioned in the methods.

When families were re-binned under adjusted thresholds (Figure 2.C, pink line) recovery of predictions improved. 29 of the expected 62 families now meet the >80% positive prediction criteria and 18 families are binned as strict positive. Demonstrating that lineage-informed thresholds effectively captures families that were excluded earlier despite conserved seed matches.

Representative cases illustrate this effect. The families that showed the highest drop out rates with the protostome cutoff: MIR-275 predictions were rescued resulting in being rebinned as strict positive, while MIR-306 and MIR-988 showed improved but only partial recovery. Some families that remained problematic were MIR-12, which continued to underperform as a result of size restrictions^23^ under general run parameters. MIR-193 and MIR-375 then paradoxically dropped in confidence and were shifted into the inaccurate bin. In the case of MIR-193 which is a microRNA strongly associated with wing color pattern variations in Lepidoptera^32^, we see that the larger proportion of predictions depict conserved seeds as expected but the non-seed bitscores were too similar to be separated efficiently.

Strikingly, MIR-219 and MIR-315 predictions failed across nearly all genomes they were found in (MIR-315 was found in 175 genomes and MIR-219 in 287 genomes). The unique predictions of MIR-219 all lacked the expected Drosophila seed motif and instead resembled an unrelated taxa, supporting its likely absence in this clade. While MIR-315, although conserved in Drosophila in its role affecting the Wg pathway^33^, was not recovered from over 80% of the lepidopteran genomes despite the cutoff adjustment. Interestingly, in the other 20% of the genomes, the top abundant MIR-315 seed predicted was found to have the conserved seed paralog as observed in the Drosophila annotation, however, the bitscores were too similar and falling short of the cutoff as is the case with MIR-193. The higher bitscores assigned by Infernal^34^ to non-seed predictions of both MIR-315 and MIR-193, could be attributed to compensatory base-pairing, as these predictions do not maintain structural similarity with forna^35^.

The adjusted threshold also results in higher cutoffs for some families leading to reduced confidence scores for those with higher numbers of expected paralogs such as MIR-3, MIR-279 and MIR-92. Families with few non-seed predictions like MIR-76 also are then more strictly binned, since the threshold is derived from a smaller pool of nonseeds, potentially exaggerating the score separation. The same is observed with MIR-275, being rebinned as strict true positive, despite some predictions falling below protostome cutoff, as there are no non-seeds to calculate an adjusted threshold from.

These results highlight that while the adjusted threshold improves confidence for many families, its reliance on the non-seed distribution assumes sufficient representation, a condition not met for some families. Thus, the adjusted threshold performs best in cases with enough non-seed predictions to allow for relative scoring and those with consistent bitscores. While not currently adapted into the automated pipeline for MirMachine 2, the adjusted thresholds serve as an effective first line approach to improve recovery in underrepresented invertebrate taxa.

### Handling missing expected microRNAs and outliers in MirMachine 2 predictions with the decision tree

Five families that remained binned as highly inaccurate despite the adjusted lineage-informed thresholds, exemplifies predictions that could not be reliably distinguished based on bitscore filtering alone. These cases illustrate a general challenge in automated microRNA annotation where bitscore based filtering alone cannot distinguish biological absence from structural divergence.

To examine the basis of this ambiguity we conducted a focused analysis on the nine economically relevant lepidopteran species (Figure 3). We therefore formalise a decision tree, where taking into consideration paralog expectations, predictions are manually evaluated using 3 main criteria - (i) Alignment to curated bonafide microRNAs of the family, (ii) presence and similarity to seed motif, especially star seeds where applicable and (iii) hairpin folding integrity, especially prolonged consecutive base pairing.

Integrating these parameters into a tree (Figure 4) style evaluation allowed us to assign the behaviour of each family to a specific underlying cause.

**Figure 4.**
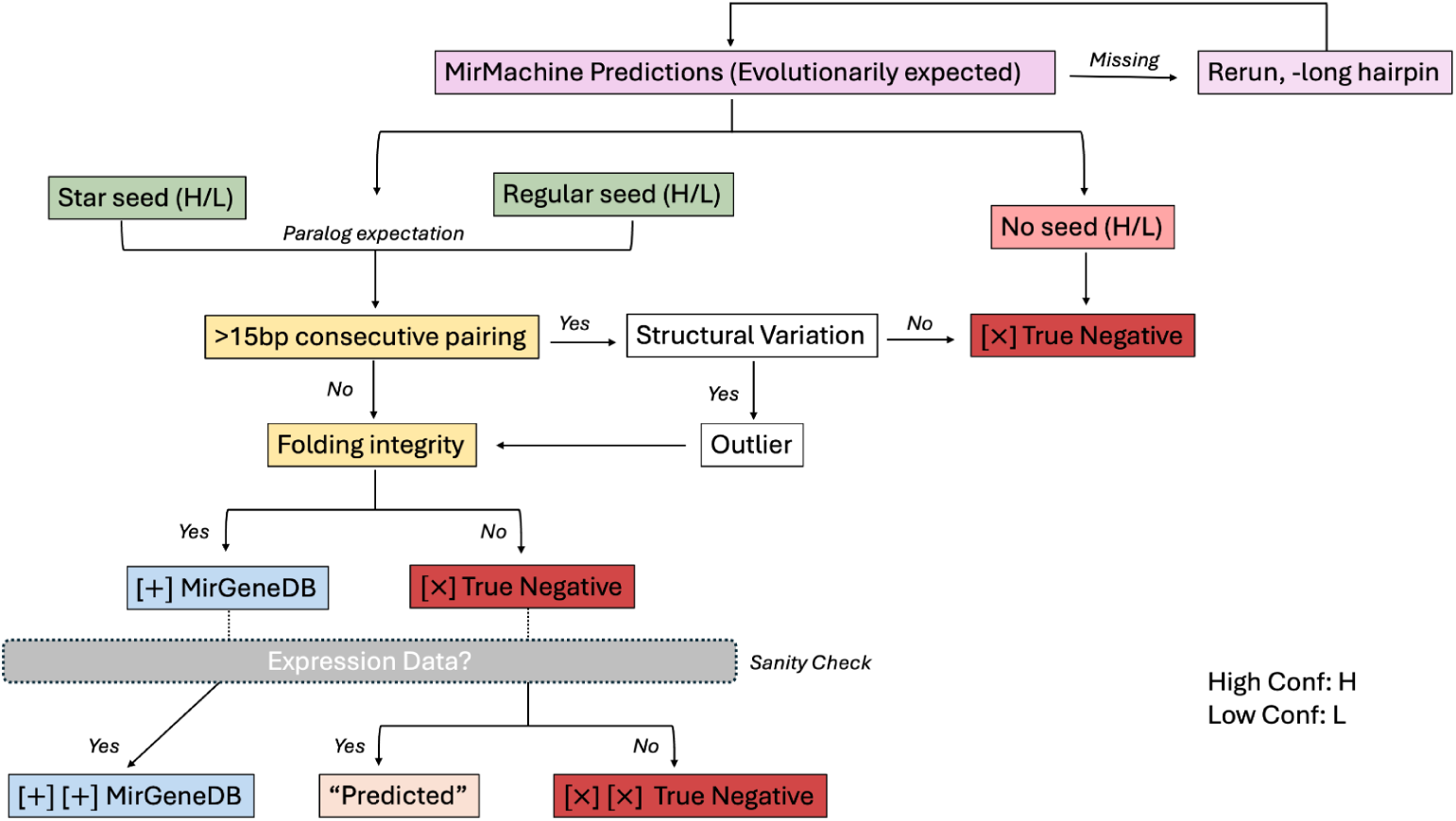
The Decision-tree framework. Workflow outlining the manual evaluation framework used to resolve ambiguous predictions. Candidate loci were assessed based on alignment, seed motif conservation and hairpin structural integrity.

#### 1. Genuine evolutionary absence

In the nine example species, all top candidates of the MIR-219 family failed sequence alignment and seed conservation criteria, confirming its absence in Lepidoptera genomes. Observed predictions of MIR-315 that retained Drosophila specific star seed motif, folded into unrealistically perfect hairpins. Applying a structural filter penalising consecutive base pairing over 15 bases, resulted in the drop out of these predictions. Thus, indicating the absence of MIR-315 as well (Supplementary Figure 3).

#### 2. Structural variability and pseudogene like remnants

Several candidates of MIR-12 displayed extended large hairpins consistent with the results reported earlier^23^ prompting the incorporation of the long hairpin parameter in addition to model refinement to accommodate broader length and structural range. This inclusion enabled the recovery of MIR-12 candidates across all 9 species, thereby improving its detection. While this option enables recovery of long hairpins that has not been achieved with existing tools, it can be time consuming. The predictions of MIR-375 and MIR-193 form 2 groups - i) Clear, high confidence, true microRNAs fulfilling all criteria ii) low scoring predictions that depict star seed conservation but aberrant folding that may represent possible pseudogenes remnants. The latter were further verified through expression data.

We applied the proposed decision tree framework across 439 lepidopteran species, selecting only the highest (N50) quality representative genome per species to assess scalability and consistency of the approach. Using seed conservation, prediction confidence and paralog expectation, we evaluated the behaviour of the five low confidence families (Figure 5). Initial results suggested the expression of the bilaterian families MIR-219 and MIR-315 which are expected to be absent in Lepidoptera. However, this signal was driven by seed representation bias skewed by predominantly deuterostome seeds. Upon incorporating lineage specific seed context, these predictions were resolved as false positives, leading to their exclusion. With the long hairpin parameter enabled, MIR-12 predictions were recovered in nearly all genomes (430), suggesting successful detection of long hairpin precursors.

**Figure 5.**
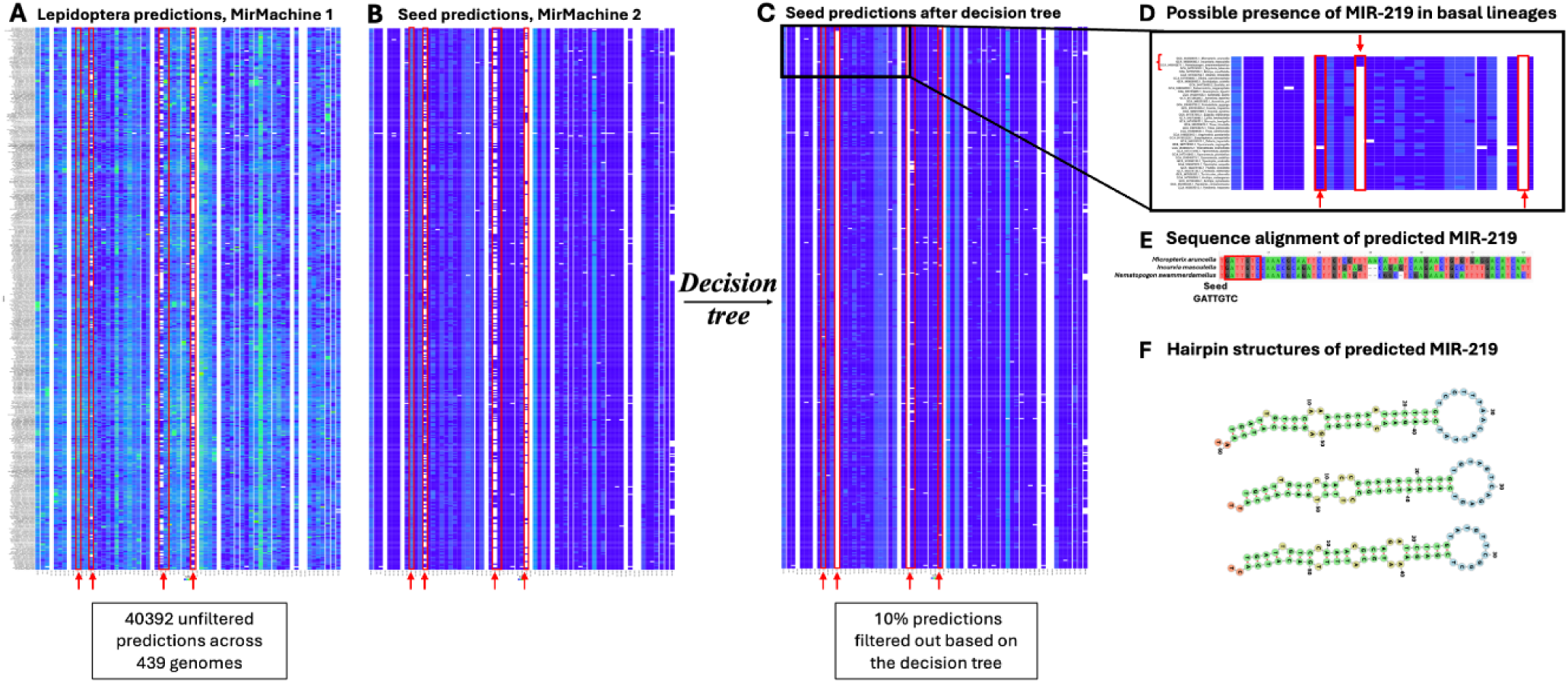
Decision tree analysis scaling across 439 genomes reveals overlooked biological signals. (A) All predictions by MirMachine 2 across high quality lepidoptera genomes. B) Predictions containing paralog informed family specific seed. C) Predictions filtered using the decision tree framework and lineage aware seed identity. D) Three basal lepidopteran species showing possible presence of MIR-219 with seed identical to Drosophila. E) Multiple sequence alignment of the high confidence predicted MIR-219 from the three genomes of basal lepidopteran species depicting GATTGTC seed conservation. F) Hairpin structures of the high confidence predicted MIR-219.

**Supplementary Figure 3.**
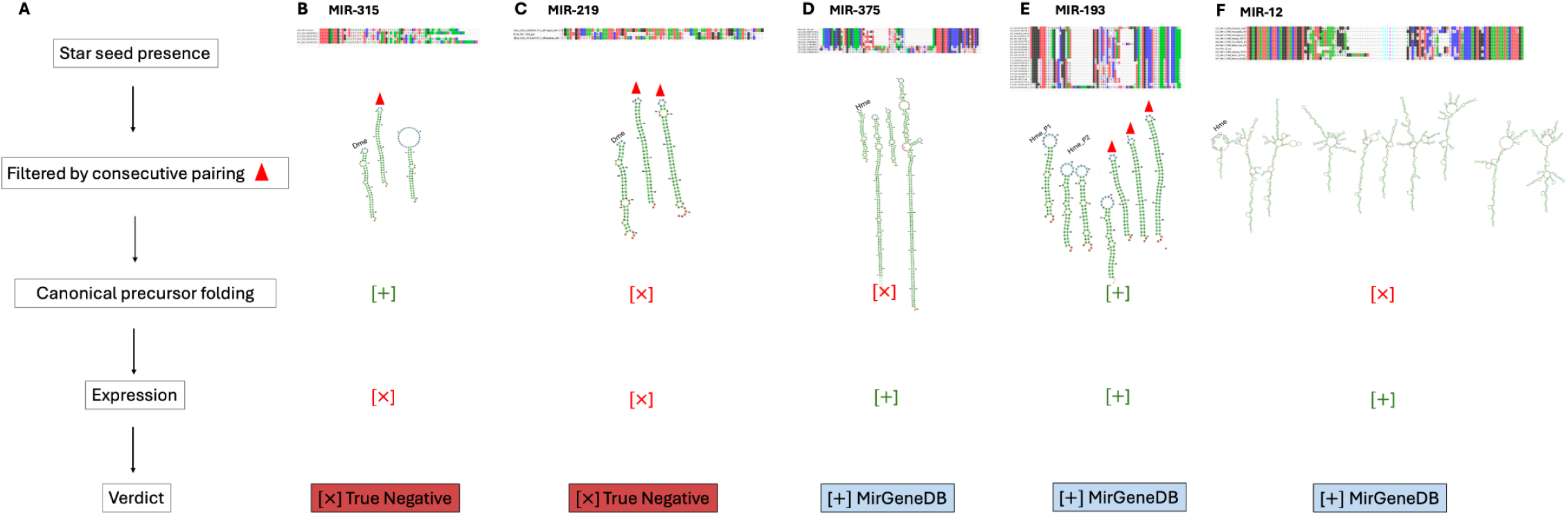
Evaluating high variability families using the decision tree. Multiple sequence alignments and secondary structure comparisons for 3 of the 5 families exhibiting high variability/persistent dropout under fixed and lineage adjusted thresholds, with MirMachine 2. For each family, selected known curated precursor structures are depicted alongside top predicted hairpins of candidate sequences. Predictions here are considered alongside the decision tree. The figure highlights cases of sequence and structural divergence that underpin annotation ambiguity. A) Simplified decision process. B) MIR-315. C) MIR-219. D) MIR-375 (structural outliers visible). E) MIR-193. F) MIR-12 (long hairpins).

Strikingly however, MIR-219 predictions persisted in three specific genomes with seed sequence identity to that of MIR-219 in Drosophila, suggesting lineage specific shared retention. In contrast, MIR-193, MIR-375 were consistent with earlier analysis. More importantly, the presence of MIR-219 in the basal lepidopteran species *Micropterix, Incurvaria* and *Nematopogon* (Figure 5.B) highlights how the updated framework enables the detection of biologically meaningful patterns that might have been otherwise obscured.

Collectively, our results indicate that while the automated approach captures the majority of conserved microRNA families, unresolved outliers warrant further interpretation. They represent biologically informative signals reflecting model limitations, structural variability and lineage specific evolutionary trajectories.

## DISCUSSION

By applying MirMachine 2 to 3019 Ensembl assemblies, we show that automated genome based annotation performs at scale without losing phylogenetic coherence. Lineage specific patterns observed across metazoan phyla represent consistent accumulation and stability of microRNAs as previously understood and are seen to be rarely lost, making them robust markers of deep evolutionary history. Through updated features including an automated microRNA score we capture aspects of genome completeness distinct from traditional coding-gene metrics. While BUSCO genes help infer genome completeness owing to widely distributed coding genes, the microRNA score derived from a smaller set of conserved non-coding elements, is sensitive to lineage specific differences in the conserved regulatory toolkit, offering a complementary perspective on the biological signal alongside genome quality with a much smaller computational footprint.

In high contiguity assemblies, microRNA scores and BUSCO converge, but systematic discrepancies arise in particular hitherto underrepresented clades, as we note in Lepidoptera. We illustrate how lineage-informed threshold adjustments substantially reduces false negatives and recovers many conserved families by adapting thresholds dynamically without full model retraining as a scalable solution. However, it exposes trade-offs causing over penalisation of families with few non-seeds and broad bitscore variance, while those with high paralog counts and indiscernible bitscores may remain obscured. These discrepancies reflect a mixture of annotation biases, family specific structural variation and in some cases genuine lineage specific losses. Distinguishing between these explanations requires orthogonal evidence. In our new version, MirMachine 2 offers conserved and star seed notations per family, to further inspect predictions suspected of underlying biological phenomenon. For example, protostome families that underperform under both fixed and adjusted thresholds like in Lepidoptera, reveal deeper biological differences rather than annotation failure, as seen with families MIR-12, MIR-375, MIR-193, MIR-315 and MIR-219. Thus, we propose a decision tree facilitating the inspection of structural integrity, sequence conservation and where necessary and possible, expression.

As observed within Lepidoptera, many apparent losses reflect methodological artefacts rather than biological signals, arising from overly conservative thresholds in underrepresented lineages, or biologically novel microRNAs (long hairpins). The detailed analyses of 9 moth genomes followed by the scaled evaluation of 439 Lepidoptera genomes, provide a practical blueprint for how targeted curation in such underrepresented lineages can improve automated annotation iteratively. Integrating lineage-aware parameters, with structured evaluation enables recovery of divergent families and improves discrimination between technical artefacts and biological signals. Families on the lower end of the confidence spectrum may require model re-training rather than threshold adjustment alone, highlighting that integrating adaptive thresholds with improved reference data will be essential.

This emphasises the requirement for systematically updated reference datasets particularly for highly diverse and genome rich clades like insects and moths (Endopterygota and Lepidoptera). To realise its full potential, we continuously integrate the expanding curated database MirGeneDB that now includes the full microRNA complements of 114 metazoan species into the model automation pipeline for MirMachine. More broadly the study reestablishes microRNAs as a practical and robust class of phylogenetic markers for large scale comparative biology. Thus, MirMachine 2 positions itself as a versatile tool for automated microRNA annotation with minimal manual interference. Particularly valuable in supporting studies of model organisms by providing a conserved microRNA profile for comparative studies and hypothesis generation in therapeutic, medical and evolutionary contexts.

In summary, MirMachine 2 resolves a fundamental limitation of genome based microRNA annotation: the lack of an evolutionarily informed, scalable and biologically interpretable scoring framework. By integrating updated and curated evolutionary knowledge, automated microRNA scoring, integration of long-hairpin models and lineage aware filtering, MirMachine 2 enables automatized and accurate representation of conserved microRNA complements and their meaningful comparison across thousands of genomes.

## LIMITATIONS

MirMachine 2 remains constrained by the taxonomic completeness and curation of the underlying reference dataset. Uneven representation in MirGeneDB can bias detection sensitivity, influence star seed estimates and contribute to apparent family absences in poorly represented taxa. Thereby limiting effectiveness of lineage informed filters and potentially misrepresenting star seed support. Although MirMachine 2 substantially improves scalability, automated annotation of thousands of genomes at scale still involves trade-offs between sensitivity and specificity. Fixed global thresholds remain overly conservative for certain lineages and while adjusted thresholds increase recovery of divergent candidates, they may increase false positives if applied inappropriately. Similarly, the new parameters, like -e value threshold and long hairpin option can increase computational demand and should be applied judiciously within the context of the decision tree. The proposed decision framework that supports targeted inspection provides a practical, but manual solution. However it introduces an element of guided curation that limits complete automation. Finally, structural and sequence based evaluation of candidate hairpins cannot substitute experimental validation of microRNA expression and function.

Addressing these limitations through expanded diverse reference datasets is the focus of ongoing work and, ultimately, must also include plant and other non-metazoan species that are currently omitted.

## STAR METHODS

Detailed methods are provided in the online version of this paper and include the following:

● KEY RESOURCES TABLE
● RESOURCE AVAILABILITY

○ Lead contact
○ Data and code availability
○ Materials availability
● METHOD DETAILS

○ MirMachine Command line tool
○ MirMachine WebApplication implementation
○ MirMachine 2 predictions
○ Retraining models
○ Runtime comparison and benchmarking
○ Binning strategy
○ Lineage aware threshold adjustment
○ Decision-tree analysis

## ACKNOWLEDGEMENTS

We thank Kevin Peterson for providing the information on evolutionary gains and losses of microRNAs, and Eric P Nawrocki who was instrumental in recalibrating the CMs for long hairpins. We are also grateful to Norwegian Research and Education Cloud (NREC) for hosting MirMachine.org.

## AUTHOR CONTRIBUTIONS

B. F and V.M.P designed the study. V.M.P performed the data analysis and generated all figures. S.U.U retrained models, updated the MirMachine software and webpage with feedback from B.F. J.A.S.T, F.F.T and L.H generated BUSCO scores and MirMachine predictions for the Ensembl dataset. V.M.P and B.F drafted the manuscript with input from all authors.

## INCLUSION AND DIVERSITY

We support inclusive, diverse and equitable conduct of research.

## STAR METHODS

KEY RESOURCES TABLE.

KEY RESOURCES TABLE:
| SOFTWARE or RESOURCE | SOURCE | IDENTIFIER |
| --- | --- | --- |
| MirGeneDB 3.0 | Clarke et al.<sup>21</sup> | https://doi.org/10.1093/nar/gkae1094 |
| MirMachine v0.3.0.3 | This study | https://pypi.org/project/MirMachine/ |
| RNAalifold 2.5.1 | Lorenz et al.<sup>36</sup> | https://doi.org/10.1186/1748-7188-6-26 |
| Infernal 1.1.4 | Nawrocki and Eddy<sup>34</sup> | https://doi.org/10.1093/bioinformatics/btt509 |
| MirMachine web app | Trondsen HT<sup>37</sup> | https://mirmachine.org/ |
| MirMachine predictions files | This study | https://github.com/VanessaPaynter/MirMachine2-resources/tree/main/Data |
| MirMachine figure data | This study | https://github.com/VanessaPaynter/MirMachine2-resources/tree/main/Figures |
| R language | R Core Team<sup>38</sup> | https://cran.r-project.org/ |
| Python | The Python Standard Library<sup>39</sup> | https://docs.python.org/3/library/index.html |

## RESOURCE AVAILABILITY

### Lead contact

Further information and requests for resources should be directed to one of the lead contacts - Vanessa M. Paynter or Bastian Fromm.

### Data and code availability

Data analyses were performed using R v4.2.0 and Python v3.9.12 custom scripts. All original code has been deposited at GitHub and is publicly available as of the date of publication. DOIs are listed in the key resources table. Any additional information required to reanalyze the data reported in this paper is available from the lead contacts upon request.

### Materials availability

This study did not generate new unique reagents.

## METHOD DETAILS

### MirMachine Command line tool

The main MirMachine engine was written in Snakemake and the command line tool, wrapper in Python v3.9.12 and R.v4.2.0 The documentation of the MirMachine CLI tool is available at our GitHub repository. It is also available as a BioConda and PyPi package for easy installation.

### MirMachine WebApplication implementation

The updated Web application is available at https://mirmachine.org. It is hosted at the Norwegian Research and Education CLoud (NREC) utilising sHPC resources.

### MirMachine 2 predictions

MirMachine 2.0 was used for the prediction of microRNAs from genomes. The Ensembl-scale predictions were generated using a custom Nextflow wrapper to orchestrate execution on the EBI Slurm cluster. For each genome, the workflow retrieved the genome FASTA, inferred the closest available MirMachine clade by comparing species lineage against NCBI taxonomy, and ran MirMachine with corresponding model configuration. Where an exact clade match was unavailable, the workflow used the nearest specific ancestral clade. The resulting prediction files were collated (Supplementary Table 2).

### Retraining models

For the model training phase of our study, we adopted the methodology previously established^22^. In short, CMs were generated for conserved microRNA families from the manually curated microRNA precursor sequences spanning 114 species from the MirGeneDB database^21^ using the Infernal RNA package^34^. For a comprehensive description of the underlying model construction and training workflow, see MirMachine publication^22^.

For MirMachine 2, we expanded the regular training workflow to build longer CMs. The loop-end position is identified from the consensus secondary structure by finding the last unpaired position before the right stem begins. At this position, the workflow adds ten identical synthetic consensus-derived sequences containing a 2000-nucleotide insert (all Ns), while the original sequences receive gaps across the same region. The modified alignment is then used to rebuild and calibrate a new CM, allowing it to tolerate long insertions in the loop region while preserving the original stem and consensus structure.

### Runtime comparison and benchmarking

A pilot dataset consisting of 71 mammal genomes (Supplementary Table 3) were used for this summary. Mean run time was used to capture overall performance across both metrics. Confidence intervals reflect variation in execution time per genome.

### Binning strategy

Taking into consideration the expected paralog number for each family, predictions containing conserved seeds were classified by comparing their bitscores to the protostome cutoffs set by MirMachine. This was used to then evaluate high scoring predictions with seed conservation, thereby narrowing true positives (Supplementary Table 4). For each family, a genome was considered to “pass” if 50% of its seed containing predictions fell within the threshold indicating a predominant number of predictions were reliably detected. Based on the proportion of genomes that met this requirement, the families were binned as follows:

- Strict true positive: 0 genomes failed, ie, all genomes show that >=50% seed-predictions for this family fall within their respective cutoffs.

- Partial false negative: Between 0 to 80% genomes failed, ie, some genomes show > 50% of seed-predictions fail to meet the family specific threshold.

- High false negative: >=80% of the genomes failed.

### Lineage aware threshold adjustment

The threshold was set at the 95th percentile of non-seed scores plus a buffer of 5 to account for potential outliers (ie, 95% of the values in the dataset are less than or equal to this value indicating most of the nonseed bitscores fall below this value and only 5% might show a higher bitscore). This ensures that the cutoff is robust, stable and not prone to extreme outliers thereby capturing typical seed signals per family.

### Decision-tree analysis

Primary analyses were conducted across nine industrially relevant lepidopteran species: *Bombyx mori* (domestic silkworm), *Grapholita molesta* (oriental fruit moth), *Spodoptera exigua* (beet army worm), *Vanessa atalanta* (red admiral), *Mamestra brassicae* (cabbage moth), P*lutella xylostella* (diamond-back moth), *Trichoplusia ni* (cabbage looper), *Manduca sexta* (tobacco horn worm), *Helicoverpa armigera* (cottonball worm).

The structured decision tree framework was applied as follows to assess consistently low scoring families:

- All unfiltered candidate predictions per genome were screened for the presence of canonical microRNA family-specific seed sequences while preserving bitscore and confidence labels.

- Predictions were then grouped by microRNA family and ranked according to descending bitscore. Within each group the top *n* candidates were selected for the next step (where *n* corresponds to the expected paralog number per family calculated from related species).

- These were then folded using RNAfold^36^, and further filtered out based on the presence of extended consecutive base pairing over 15bases in the hairpin structure.

- In cases where predictions of expected conserved families were failing the previous step, and thus were not recovered (as seen in the case of MIR-12 among the nine lepidopteran species), genomes were reprocessed using the long hairpin parameter. The same MIR-12 targeted pipeline was subsequently applied to the 439 lepidopteran genomes (Supplementary Table 5), selected on the basis of high N50 enabling comparative evaluation to scale.

